# Three-dimensional Tracking Method for Water-Hopping Mudskippers in Natural Habitats

**DOI:** 10.1101/2025.04.02.646728

**Authors:** Daehyun Choi, Kai Yung, Ian Bergerson, Halley Wallace, Ulmar Grafe, Saad Bhamla

## Abstract

We present a portable, non-invasive, and low-cost three-dimensional tracking method to quantify in situ water-hopping kinematics of mudskippers. By combining dual-camera video recordings with tracking fish path, Gaussian Splatting terrain reconstruction and epipolar geometric analysis, we capture detailed 3D trajectories of mudskippers in their natural tidal-flat habitats. Our proposed method resolves complex hopping motions, including both straight and curved escape paths, and reveals that horizontal distance, hopping height, and speed are strongly influenced by fish size and local terrain features. These results highlight both the biomechanical and ecological significance of water-hopping in mudskippers, demonstrating how a simple, deployable 3D approach can resolve complex amphibious movements in challenging field environments.

## Introduction

Mudskippers are uniquely adapted amphibious fish capable of dynamic terrestrial and aquatic locomotion (Figure 1a-b), making them an ideal subject for studying the evolution of land-based movement in vertebrates. Their ability to move on land has been extensively studied [1, 2, 3], yet data on hopping behavior over water remain sparse (Figure 1c). Although a few studies have mentioned this behavior in laboratory conditions [4] and in situ [5], a comprehensive analysis of their three-dimensional kinematics while interacting with their heterogeneous environment has not, to our knowledge, yet been performed. Early documentation of mudskipper locomotion can be traced back to Petit (1921) [6], who noted that each bound is preceded by a brief period of swimming, describing it as “ricochet”. Harris [7] later termed the behavior “skimming,” emphasizing that the mudskipper actively powers its series of hops across the water using its caudal fin, rather than its hops being a passive bounce like a skipping stone, which is propelled by its spin across the surface of water [8, 9]. Harris described the skimming sequence in detail: (1) the fish swims normally with its eyes and part of its head above water, (2) it accelerates to exit the water at about a 30 angle (with respect to the water surface), (3) it remains airborne, and (4) it lands back on the water surface. Harris reported an average velocity of 2.5, m s^−1^ for a 14 cm body length, corresponding to 17.9, BL s^−1^ (where BL denotes body length). Based on schematic observations, Harris suggested that the mudskipper does not leverage its pectoral fins. However, modern imaging reveals that pectoral fins are actively used and are crucial for skimming dynamics [5] [10]. From this point on, we describe this behavior as “hopping”, and note the mudskipper has different methods to control the rate of hopping and how often it emerges from water. Our goal is to quantitatively analyze the hopping behavior using multiple cameras to provide a fuller description of mudskipper hopping behavior in the field.

**Fig. 1.**
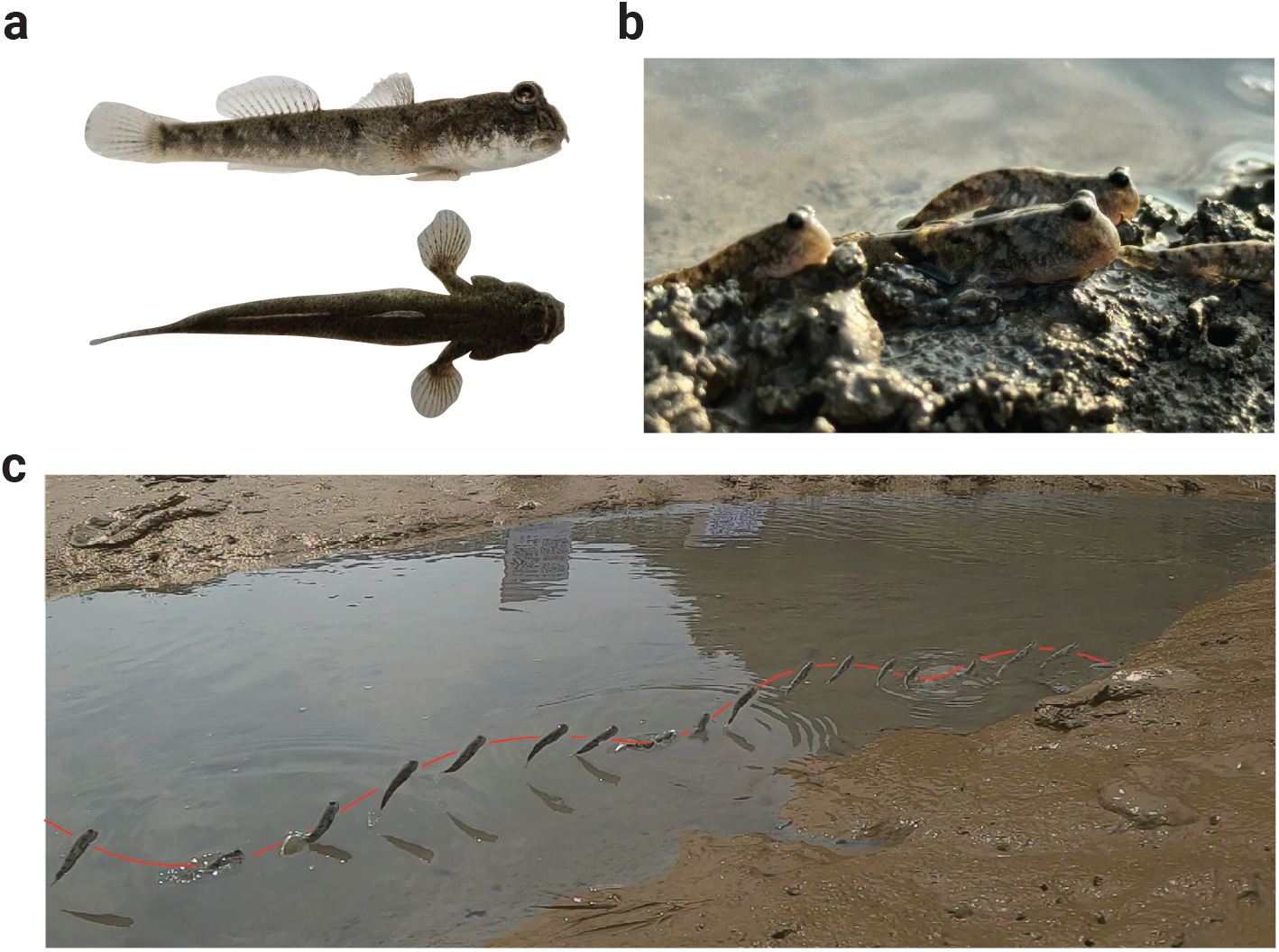
*Periophthalmus modestus* in natural habitat. (**A**) Side and top view of live mudskipper. The pectoral fin is retracted in side view while extended in top view. (**B**) Mudskippers resting on the bank. (**C**) Time steps of a progression of a mudskipper hopping across the water’s surface.

Although this behavior is often featured in popular media (e.g., Netflix documentaries and Seoul Broadcasting System), Wicaksono et al. (2020) [5] provided one of the first detailed kinematics report of mudskipper locomotion, classifying mudskipper hopping as a novel form of fish locomotion distinct from water-jumping or the gliding of flying fish. By using an action camera, they documented hopping, taxiing, submergence, and combinations thereof on land, water surfaces, and tree roots, reporting velocities, time durations, and energy-loss analyses. However, using a single-camera approach lacks in-plane motion information. While a two-camera setup to capture a 3D recording can be more accurate, it is more convenient on flat surfaces where using epipolar geometry, lines from both camera views align to identify the same pixel pattern, and such ideal conditions rarely exist in natural habitats. In other organisms, methods of air-water interface locomotion, such as skimming frogs [11], or basilisk lizards [12] can be analyzed in lab with 3D set ups, but acquiring full 3D trajectory data is often cost-prohibitive and challenging for animals sensitive to external disturbances, particularly in situ for aquatic animals living in dynamic intertidal zones with continually changing water levels.

Recently, advances in 3D reconstruction algorithms (e.g., Gaussian Splatting, LiDAR) have facilitated the use of low-cost sensors (e.g., cell phones, webcams) for capturing high-resolution data. Vieira et al. (2025) [13], for example, achieved high spatio-temporal resolution measurements of ocean waves using two low-cost cameras. Inspired by this, we introduce a novel, low-cost, and highly portable 3D tracking method for natural habitats, which combines an existing 3D Gaussian Splatting algorithm (using the commercial software PolyCam [14, 15]) with 3D epipolar line reconstruction using in-house code (Matlab).

This study provides two main contributions. First, we detail our field-ready 3D imaging technique that allows unobtrusive monitoring of amphibious fish in their native environment. Second, we demonstrate that this method yields high fidelity motion trajectories, enabling high spatio-temporal resolution analysis with a spatial resolution of 0.419mm near the center of the field of view (FoV) (which may vary depending on camera configuration, pixel location, lens distortion, and epipolar angle relative to the centerline) and a temporal resolution of 4.17ms, for analyzing hopping performance and terrestrial navigation within a wide FoV of approximately 1.5 *×* 1.5 *×* 1.0m in the *x, y* (horizontal), and *z* (gravity) directions, respectively. Overall, our results highlight the efficacy of non-invasive, three-dimensional tracking as a powerful tool for studying the biomechanics and ecology of amphibious animals in their natural environments.

## Method

### Location Selection

We selected two tidal flat areas on two islands (Daebudo in South Korea and Bedukang in Brunei) for mudskipper observation. Mudskippers mainly inhabit brackish areas where seawater and freshwater meet, where Ikebe and Oishi observed the same species in Figure 1 (*Periophthalmus modestus*) while studying their behavior during high tide [16] and documented earlier by Macnae (1968) [17]. After identifying a few possible mudskipper sighting locations through internet research, we conducted a preliminary site survey and interviews with local residents 1-2 days before the observation date. We selected these locations because there was a sufficient concentration of mudskippers (Figure 1b) and also considered the level of safety and accessibility. Daebudo Island is located near private land right next to a road, and in Brunei we hired a captain to reach the uninhabited island of Bedukang. For reference, the Brunei trip was pre-approved by the authorities (approval number : UBD/AVC-RI/1.21.1[a]/2024/009). The coordinates are provided in the figure captions for the specific locations.

The observation area was located where freshwater flows into the sea (estuarine zone, see Figure 2b). During our fieldwork, this area had the highest density of mudskippers and showed frequent hopping behavior. Another factor in choosing this area was its easy accessibility for people. However, due to the tidal characteristics, the sea level changes over time (Figures 2a and 2c) and this area becomes completely submerged during high tide (Figure 2d). Therefore, we limited the maximum observation time to three hours and developed a methodology to complete all experiments within that period.

**Fig. 2.**
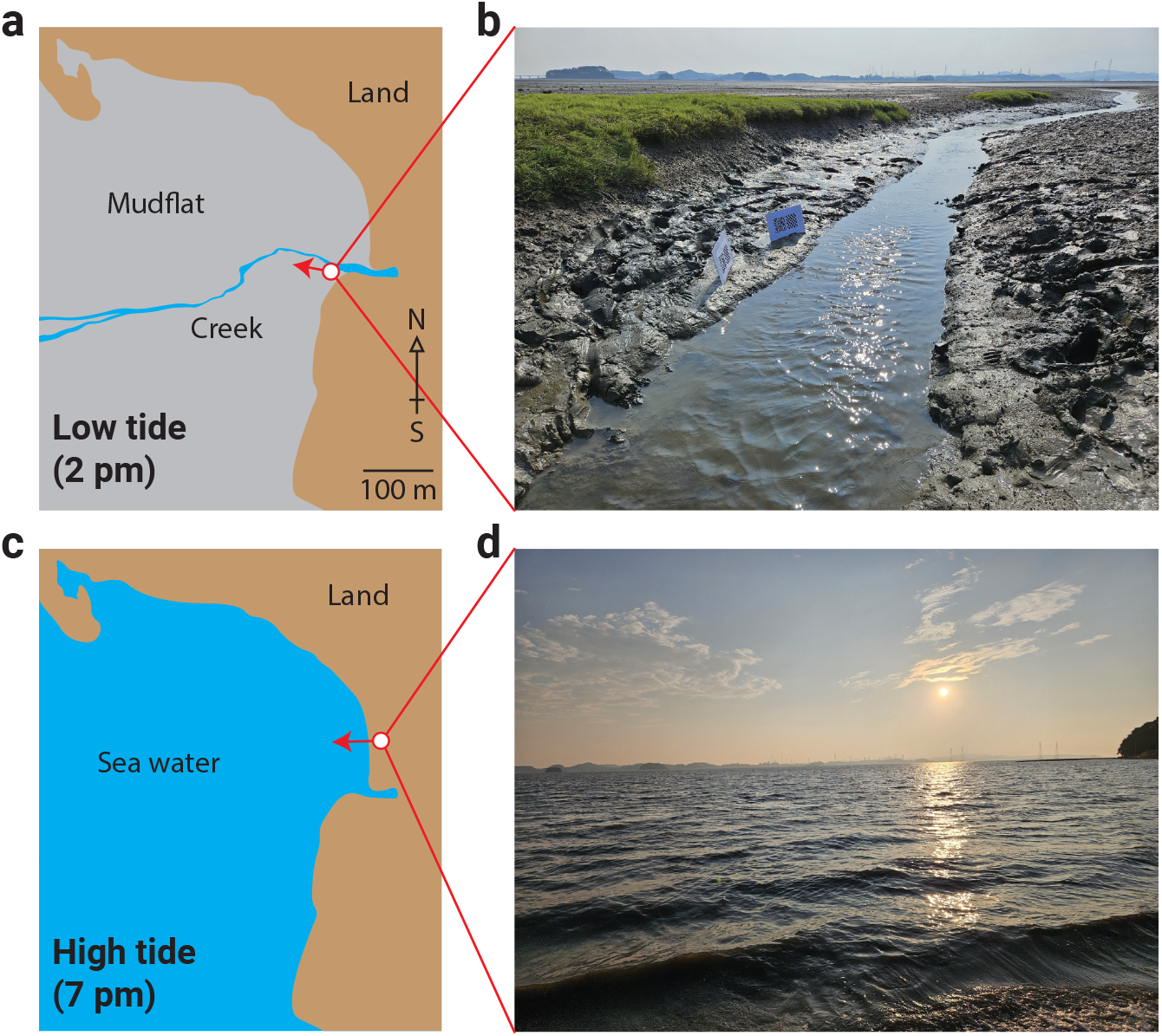
Natural habitat of mudflat area in Daebudo, South Korea. **(A-B)** High tide; **(C-D)** Low tide. The arrows in **A** and **C** denote the direction of the camera which captured **B** and **D**, respectively. See acknowledgment for detailed location and date.

### Camera Configuration and Measurement Setup

The locomotion of the mudskippers on site was captured by two portable cameras (GoPro Hero 12) positioned at different locations (Figure 3a). Each camera’s resolution was set to 2704 × 1520 pixels (2.7k) at 240 fps, with an approximate viewing angle of 120°. Both cameras were oriented toward the area where the mudskipper hoppings occur and were fixed on tripods such that their lines of sight formed an angle of roughly 90° with each other, while the angle varied depending on the terrain and environment. Two calibration targets were placed on the opposite side of the cameras (Figure 3b) to allow for calibrating length and camera orientation.

**Fig. 3.**
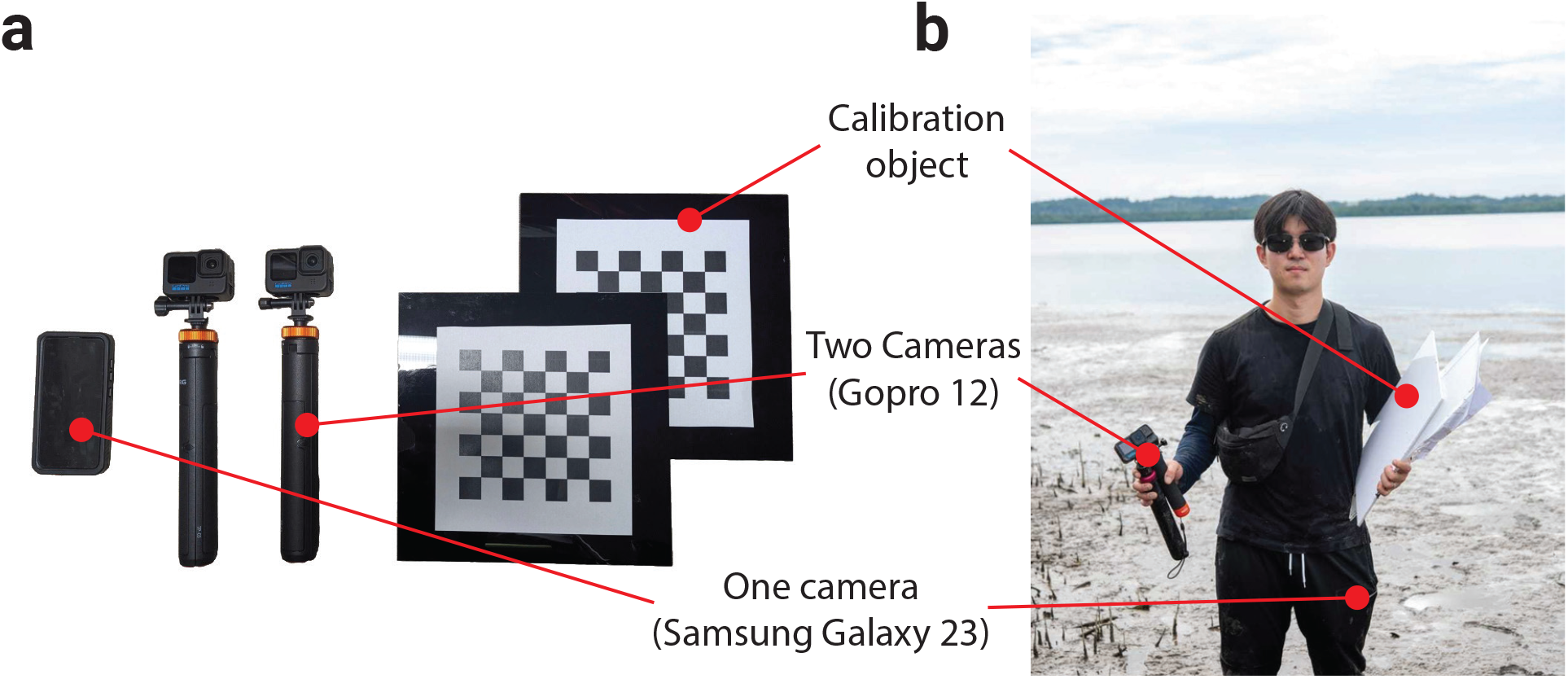
(**A**) Experiment equipment (Two action cameras (GOPRO) and tripods, one cell phone, and two calibrations targets), (**B**) Setup demonstration, and (**C**) Carrying demonstration in field (Pulau Bedukang). Image includes the first author, with permission granted for publication.

### Terrain Reconstruction Using Gaussian Splatting

To obtain three-dimensional terrain information, multiple snapshots were taken with a high-resolution camera (Samsung Galaxy 21) from various angles (Figure 4b). Using these images, a 3D point cloud was constructed with the Gaussian splatting algorithm (PolyCam - LIDAR & 3D Scanner [14]) (Figure 4c). Since this data not only contains the terrain information but also the camera positions, there is a benefit in being able to select arbitrary camera locations during experiments while still acquiring accurate relative positions of the calibration patterns in three-dimensional space. Accordingly, one can place a virtual camera in the virtual terrain at the same location as the experimental camera positions, then, utilizing the calibration and terrain data, the camera angle and focal length parameters can be fine-tuned to replicate the real-world camera view (compare the left and right sides of Figure 4d and 4e). Here, Figures 4d and 4e show photographs taken by two cameras, with the mudskipper’s trajectory indicated by a dotted line and its instantaneous positions marked by red boxes. The mudskipper traverses a complex three-dimensional path while moving to different locations on the land.

**Fig. 4.**
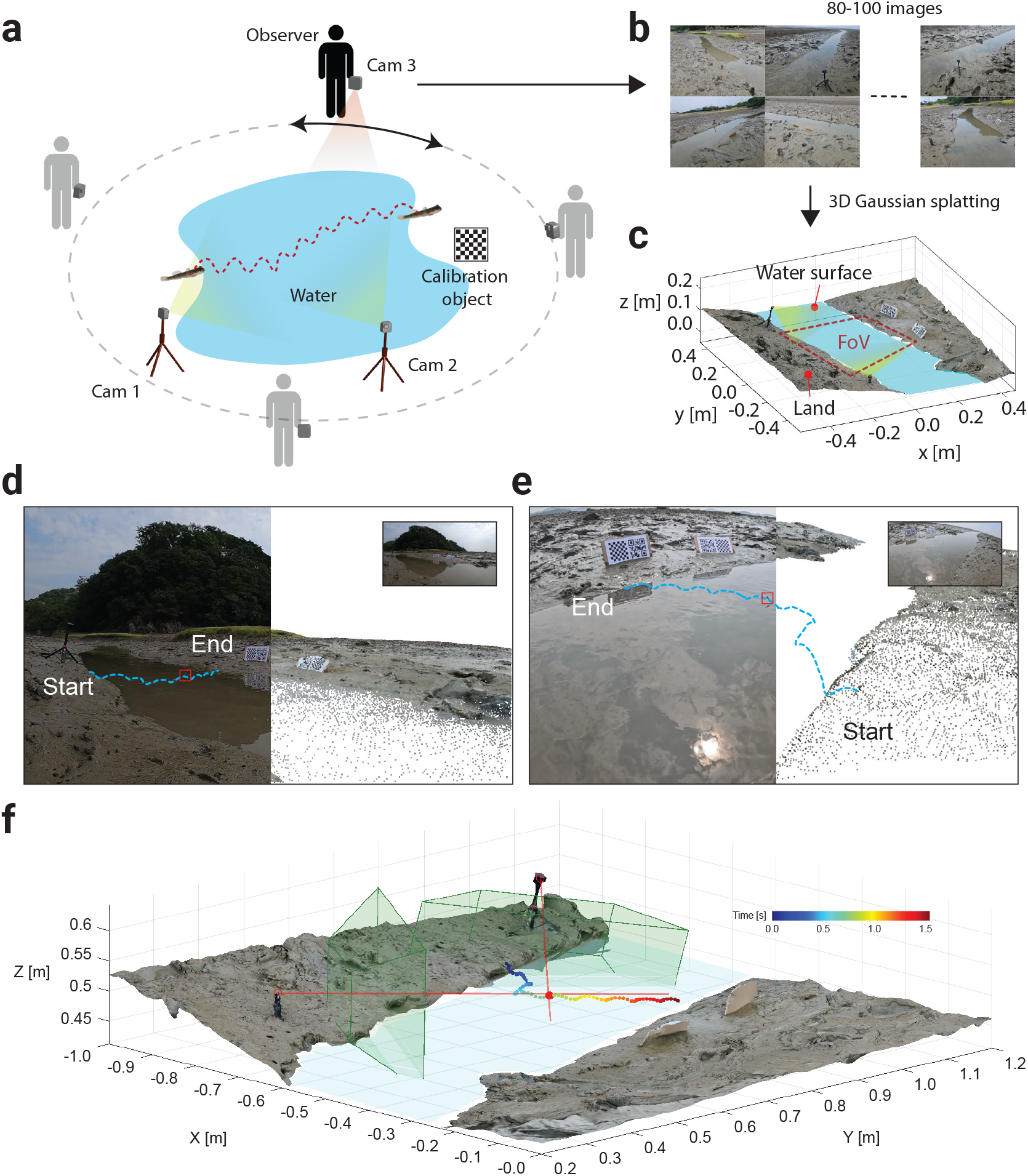
Method to obtain the 3D trajectory of mudskipper. (**A**) Measurement setup. (**B**) Raw image of snapshots for local terrain. (**C**) The reconstructed 3D terrain from 3D Gaussian splatting. (**D-E**) Camera view (left part) and the virtual camera view (right part) from left (**D**) and right camera (**E**). (**F**) The reconstructed 3D trajectory by 3D ray tracing from both camera. Here, camera intrinsics such as focal depth and focal angle is visualized as frustums.

### Experimental Procedure and Data Acquisition

The experimental steps were as follows:

1. First, we selected a mudskipper-dense region. In this region, we commonly found the mudskipper species *Periophthalmus modestus* and *Periophthalmus gracilis*.
2. We placed Cam1, Cam2, and the calibration targets to allow clear observation of this region, and recording was started (5 minutes required).
3. Next, we took terrain snapshots from multiple angles using the Smartphone camera (another 5 minutes).
4. During the same 15-minute interval, Cam1 and Cam2 recorded continuously while the camera operator waved a long stick or hand from a distance to provoke the mudskippers to perform escape behavior (without making direct contact).
5. After recording, we removed the equipment and repeated the experiment at two more locations, for a total of three trials. We collected sufficient data during the high-tide period.

### 3D Trajectory Extraction

To derive three-dimensional position data of the mudskipper trajectories, the time-series trajectories were first tracked in each camera view (using DLTdv8 in MATLAB [18]) to obtain their pixel coordinates. The sampling interval between trajectory points was 5 frames (i.e., 0.021 seconds, given the 240 Hz frame rate), and a total of *n* = 48 trajectories were extracted.

### Epipolar Geometry and 3D Position Estimation

By applying the known (and fine-tuned using 3d point cloud data) focal length, camera positions, and angles, each trajectory point could be mapped onto its corresponding epipolar line in 3D space (Figure 4f). The intersection of the two epipolar lines from the two cameras indicates the mudskipper’s three-dimensional location. However, because of small errors in camera parameters and tracking data, the two lines did not intersect perfectly. Therefore, the mudskipper’s 3D position was approximated by taking the midpoint of the closest pair of points on the two skew lines. All calculations, including editing the 3D terrain data, generating epipolar lines, and computing intersection points were performed in MATLAB using custom, in-house code. These scripts have been shared on GitHub (github.com/bhamla-lab/mudskipper-3D-tracking-ICB-2025) for reproducibility.

## Results

### Representative trajectories

Figures 5a–c present three representative three-dimensional trajectories of mudskippers, demonstrating three typical methods of locomotion that mudskippers took. By employing this 3D measurement technique, we quantitatively captured both the in-plane and out-of-plane movements of the mudskippers in detail. In both Periodic hopping 1 and 2, the mudskippers display periodic hopping. In Periodic hopping 1, if we neglect the motion in the z-direction, the mudskipper moves in a straight line (small curvature) and reaches the opposite shore, whereas in Periodic hopping 2, it follows a large curvature to arrive back at the same shore. Figures 5c shows the instantaneous velocity with fish schematics, indicating that the speed for the trajectory peaks at each hopping and decelerates when the fish meets the water surface. A comparison of average speed and hopping performance as a function of curvature is discussed in more detail later in the statistical analysis.

**Fig. 5.**
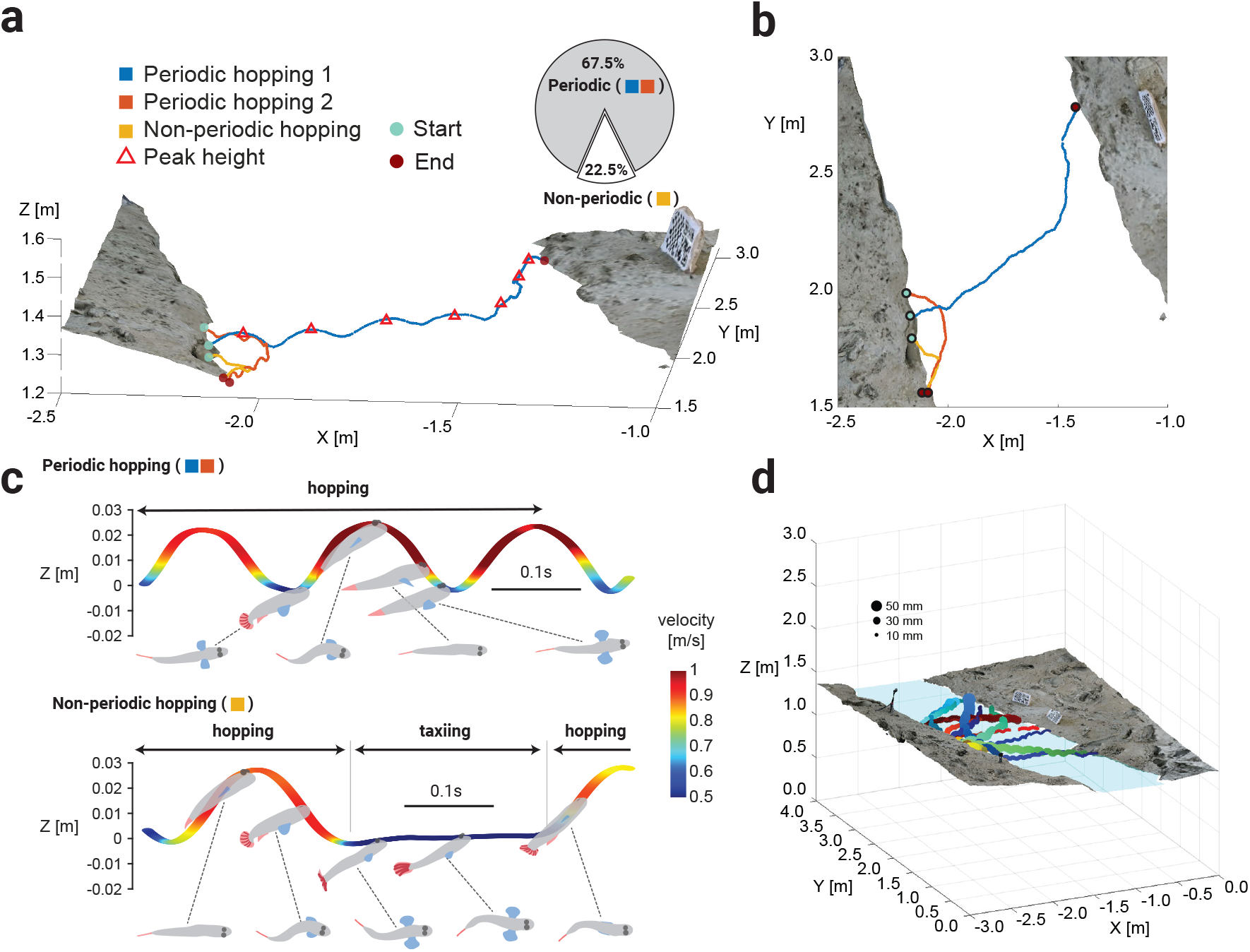
(**A**) Three representative trajectories of mudskipper paths tracked on the 3D plane, where ‘Periodic hopping 1’ traces a mudskipper traveling across the water from one piece of land to another, the ‘Periodic hopping 2’ and ‘Non-periodic hopping’ show mudskippers which traveled onto water but returned back to their starting piece of land. In ‘Non-periodic hopping’, the mudskipper taxied intermittently between hops. The pie chart shows the occurrence rates are 87.5% and 12.5% (n=48) for periodic and non-periodic trajectories, respectively. (**B**) Top view of the three trajectories. (**C**) Two hopping mode with schematics showing fins and body orientations: Top and bottom view correspond to periodic and non-periodic hopping, respectively. (**D**) Multiple trajectories of mudskipper paths in 3D space for one site.

Non-periodic hopping involves movement that is not solely consecutive hopping; rather, it includes “taxiing” in the intervals or before/after hopping. Taxiing differs from typical underwater swimming in that part of the fish’s eyes and head remain exposed to air during movement (see schematics for taxiing in Figure 5c). Its fin usage is also distinctive: the pectoral fins remain extended to generate lift, while the caudal fin produces thrust. In Non-periodic hopping, the fish jumps from land to water and then continues taxiing for around 0.2 s with extended pectoral fins—leading to high drag and consequently low speed (*<* 0.5*m/s*). It subsequently returns to the same land by hopping. Such trajectories that mix hopping and taxiing are found in both trajectories that return to the same shore and trajectories where the mudskipper crosses the water to another shore. We classify these irregularly mixed hopping behaviors as “non-periodic hopping.”

Hence, based on our observations, mudskipper trajectories can be broadly categorized into two types, periodic and non-periodic trajectories, whose occurrence rates are 87.5% and 12.5% (n=48), respectively (see pie chart in Figure5ca), indicating the fish prefers the periodic hopping. Example videos of periodic and non-periodic trajectories are found in SI video 1. By determining the 3D positions of the fish’s head and tail, this technique (Figure 4f) enables us to estimate the fish’s size. For *P. modestus*, the total length (TL) is 24.6 *±* 10.8*mm*) (n=41), while for *P. gracilis*, 54.5 *±* 2.9*mm* (n=2). Baeck et al. [19] reported *P. modestus* in the range of 31–99 mm (TL), which is higher than our measurements, indicating that the mudskippers in our current observations are mostly premature. The length of *P. gracilis* ranges from 25–40 mm in standard length (SL) [20], and its maximum length is comparable to our measurements (noting that our values are in total length), suggesting that the observed individuals are fully grown.

### Statistics analysis

Figures 6a–b illustrate the average hopping speed and total path length of the trajectory as functions of fish body length. In general, the average speed and distance for escape behaviors generally increases with body length [21, 22]. The mudskippers observed here also exhibit proportionality between speed and body length, but not distance, which might result from the geometry-sensitive nature of water-hopping locomotion. This could be due to the irregular shape of water surface or interruptions caused by land features and dense mangrove roots, which often led to early termination of hopping. This aspect should be refined in future studies. In this study, the maximum speed normalized by body length reached 70 BL/s (where BL is body length), and the distance covered in a single trajectory was 125 BL.

**Fig. 6.**
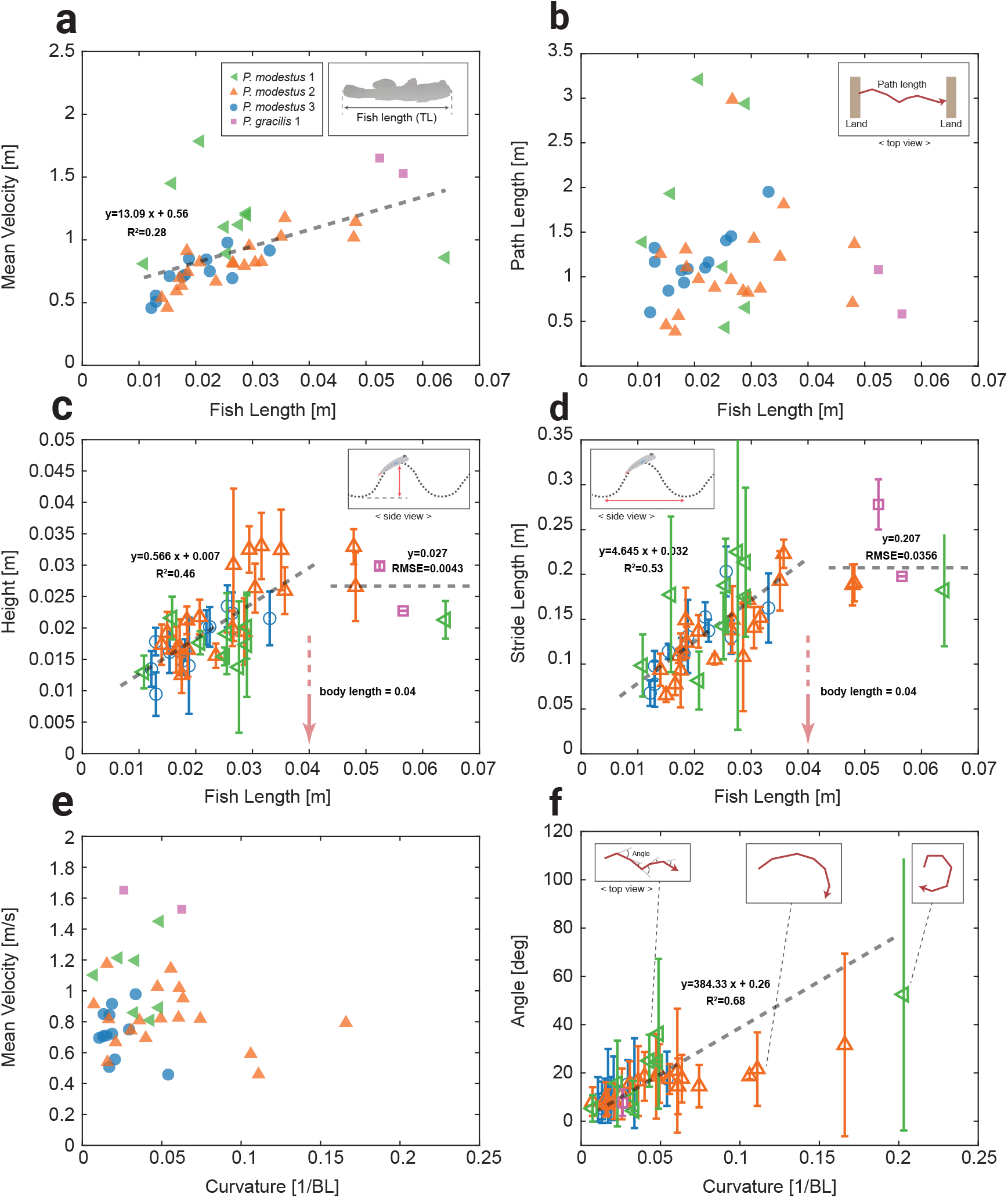
Statistical features of hopping behavior collected from 3D trajectories. (**A**) Mean velocity, (**B**) Path length, (**C**) Height, and (**D**) Stride length as functions of fish length. (**E**) Mean velocity and (**F**) Turning angle *versus* maximum curvature along the path. Each plot includes 42 trajectories: 40 from South Korea (*P. modestus*) and 2 from Brunei (*P. gracilis*). Error bars indicate standard deviation.

Next, Figures 6c–d show the horizontal and vertical hopping distances for individual hopping segments. A single trajectory can contain multiple hopping segments, and each symbol in these figures denotes the mean and variance of the parameters for all hopping segments in one trajectory (corresponding to one fish length). The horizontal and vertical distances increase linearly with fish body length, but stop increasing and remain constant beyond 0.04 m (see the arrows in figures 6c and 6d). When the fish’s body length is below 0.04 m, it can jump up to its own body length in height. Horizontally, the mudskipper can leap to about five times its own body length (when below the critical length). For mudskippers larger than the critical length, both vertical and horizontal leap distances tend to remain constant—approximately 0.027 m and 0.21 m, respectively. Additional observations of mudskippers exceeding the critical body length are needed, necessitating further field work in the species’ native habitat. While the fluid dynamic rationale for designating 0.04 m as the critical length warrants investigation, it lies outside the scope of this study and will be pursued in future work.

Lastly, Figures 6e–f present the average speed as a function of curvature and the change in horizontal angle (in the z-plane) between hopping segments. From Figure 6f, one can see that the average speed is hardly affected by small curvature below 0.1 [1/BL] after which mean velocity is significantly decreased *<* 1*m/s*. We can infer that for more curved trajectories (*>* 0.1 [1/BL]), part of the horizontal momentum is lost before the next leap, and assuming the same energy expenditure, this leads to lower overall speed. In addition, it appears that mudskippers can adjust the horizontal angle between each hopping segment in proportion to the curvature (Figures 6f).

## Discussion

### Comparisons with Other Amphibious Organisms

The mudskipper kinematics observed here, in contrast to data from laboratory mudskippers [4] and amphibious animals like frogs [11] or the plumed basilisk lizard [23], can help elucidate the limits of water-surface traversal mechanisms. Weiss [11] indicates that the skittering frog follows a primarily ballistic trajectory, reaching a maximum velocity of approximately 1.1 m/s, and the basilisk is reported to run at about 1.3 m/s [23]. These velocities are similar to the mudskippers mean velocity, signifying that water-surface traversal methods may have a maximum limit between 1 and 2 m/s. This claim is further reinforced by the water strider, *Gerridae*, which is reported to have peak speeds of 1.5 m/s as well [24]. SVL length scales of the basilisk and skittering frog are on the order of 0.1 [25] [11], while water striders and mudskipppers (studied here) are on the order of 0.01 [24] so this limit is known to apply for a small range sizes. The exact reason behind this suggested limit is currently unknown, but hydrodynamic drag or biomechanical limitations may be the primary reason why animals on the water surface can’t break this speed limit. Other animals, like the flying fish *Exocoetidae*, are seen breaking this limit at approximately 15–20 m/s [26], although they don’t need to follow this rule. The important distinction to make between these scenarios is that the flying fish leaves the surface of the water. Since it is not bound to the surface in the same way as the other examples, it can exceed the limit. Once it does return to the surface, it breaks the surface tension and returns below.

### Convergent Evolution in Water-Surface Hopping

An interesting comparison involves the Chinese Rice Grasshopper, *Oxya chinensis*. Although not closely related to the mudskipper, it exhibits a jumping behavior in water similar to that of the mudskipper, as reported by Song et al. [27]. Both species use a caudal structure to propel themselves out of the water: the grasshopper uses its hind legs, and the mudskipper its caudal fin. In both cases, this mechanism functions as an escape from predators. This similarity can be seen as a convergent evolution, and deeper investigations may have significant implications for bio-inspired designs featuring adaptable, multimodal locomotion. Such designs could address diverse environmental scenarios, including muddy, wet, and uneven surfaces (as seen in the mudskipper), flight (as in the grasshopper), and the common link of hopping to escape via water.

### Evidence for Ballistic Motion

Our imaging analysis indicates that the mudskipper’s parabolic hops resemble the ballistic motion observed in the African Butterfly Fish [28] and the skittering frog [11], albeit with a distinct take-off mechanism. The African Butterfly Fish uses an abduction of its fins to propel itself upward [28], whereas the skittering frog does a standard jump but in the water[11], and the mudskipper launches itself by rapidly straightening its caudal fin, previously bent into a C-shape [5]. Despite differing modes of propulsion, all follow a similarly parabolic path.

Saidel et al. [28] identify several criteria characterizing ballistic motion in the African Butterfly Fish; our findings align with five of their points:

1. The mudskipper’s short take-off period occurs when the fish bends its caudal fin and quickly extends it.
2. A single peak in the path is evident in the trajectory data.
3. Stride length spans a few body lengths (as shown in Figures 6d).
4. Velocities remain below those of flying fish *Exocoetidae* (15–20 m/s) [26].
5. The observed path is parabolic.

There is insufficient data to confirm the sixth criterion of short air time compared to flying fish. However, the parabolic arc suggests no significant gliding: the mudskipper’s time aloft is notably shorter than flying fish, indicating ballistic motion, rather than sustained flight.

### Directional Preference as an Escape Mechanism

The high frequency of smaller curvatures, corresponding to turning angles *<* 40^°^, suggests that mudskippers typically move in a single direction with minimal veering. This may be because altering the course could lead back toward the original danger, introduce new danger, or require more energy. Consequently, this locomotion is consistent with an escape mechanism, as noted by Harris [7], although Davenport observed that flying fish sometimes employ flying to reach food sources [26]. Overall, water hopping in mudskippers operates as a multifunctional traversal method, yet its out-of-water motion follows a predominantly ballistic trajectory, distinct from the sustained flight of other species.

### Limitation and Application of the Proposed Method

This method requires four major constraints.

1. The camera position must remain fixed during recording. It cannot be placed on any surface that moves, such as a vessel in motion or a swaying tree branch.
2. The observer must have sufficient accessibility to collect enough snapshots of the terrain on which the organism moves. In this study, mudskippers lived in a creek that reached only thigh depth or in puddles entirely surrounded by land, allowing multiple-angle snapshots. If such viewpoints are not possible, other measures need to be explored.
3. The target organisms must stay within or return to the field of view after setup and snapshot acquisition. Although the mudskippers tended to move away from observers due to their sensitivity, their very high density ensured they were always present in the observation area. Otherwise, one would need either less invasive methods for snapshots or other techniques for terrain measurement.
4. Organisms must be optically visible. Movements become difficult to measure if they frequently dive or hide in burrows. Mudskippers, however, expose much of their eyes and head to air while hopping, so this method remains feasible.

When these conditions are met, the proposed approach offers high temporal resolution (e.g., 4.2 ms in this study) and high spatial resolution (e.g., 0.5 mm) over a broad observation domain (e.g., 1.5 *×* 1.5 *×* 1 m). For instance, water striders approximately 50 mm in size, found in the same habitat as

*P. modestus*, were also observed hopping across the water surface and traveling distances exceeding 2 m at speeds of about 1 m*/*s. Their three-dimensional movement could be adequately captured with this technique. This approach is not limited to centimeter-scale organisms. By rearranging the camera placement, for instance on exposed rocks at the ocean’s edge, it becomes feasible to measure three-dimensional movements over tens of meters, including sea otters swimming at the water surface or pelicans hunting while floating, as well as the relationships between these animals and surrounding terrain. Terrestrial and aerial organisms, such as rabbits moving on land or insects flying and hopping, could also be observed provided the terrain is captured from multiple angles in the limited time. This method is particularly effective in situations where the observer must operate in random and unpredictable terrain, or when filming needs to be completed within a short time frame.

It is necessary to discuss potential errors in organism positioning caused by inaccuracies in the camera’s intrinsic and extrinsic parameters. This method adjusts the camera’s position, orientation, and focal length by manually matching the real camera view with the reconstructed terrain’s virtual camera view. Future refinements using advanced techniques, such as cross-correlation based on light intensity and color information in the reconstructed terrain, or methods like Shake-the-Box, could further automate and enhance this calibration process, which is expected to improve the overall methodology.

## Conclusion

We developed a method for capturing the 3D trajectories of mudskippers in situ were able to measure the hopping paths of *Periophthalmus modestus* and *Periophthalmus gracilis* in the intertidal regions of Daebudo Island, South Korea, and Pulau Bedukang Island, Brunei. The recorded trajectories gave a way to find the velocities at each point along the trajectory, as well as the turning angle from the initial direction. The measurements revealed that the mudskippers experience ballistic motion, shown by parabolic arcs in the recorded hopping motion. The heights and stride lengths of those arcs were observed to increase with the body length of the mudskipper, but the total path length did not. Most importantly, the distance traveled did not affect the mean velocity the mudskippers would hop at, which could show that the mudskipper may use this locomotion technique as an efficient form of travel instead of just as an escape mechanism [7].

There are many more mudskipper species that this imaging and tracking technique can be applied to, which could reinforce the findings of *Periophthalmus modestus* here or indicate that species have varying hopping characteristics. Furthermore, analyzing other species of mudskipper may lead to a better understanding of the ecological importance and the behavioral characteristics of their skipping behavior, it could also give insight into how the skipping behavior evolved across species. Applications in other systems can lead to new discoveries in locomotion behaviors in the context of their environment. Future studies using this method could introduce the ability to track individual organisms across different media, a capability lacking in this set up. Currently, vision of mudskippers is lost when they fully submerge after a hop, but a novel method of recording their underwater motion could be developed to counteract this limitation. Mudskippers aside, it can also be used to collect in situ data for a variety of biological organisms which is very difficult to obtain. By collecting data which more accurately represents how organisms locomote in their environment, future bio-inspired robotic designs can better represent their model organism, possibly leading to improved functionality. Thus, this method of recording mudskippers in situ is an effective and non-invasive way to gain data about mudskipper hopping trajectories and allows us to learn more about their water hopping behavior. Additionally, this method can be easily applied to many different biological systems because of its low cost and flexible set-up.

## Author contributions statement

D.C. conceived, conducted experiments, analyzed the results, wrote, and reviewed the manuscript. K.Y. obtained, anaylzed tracking data, wrote, and reviewed the manuscript. H.W. obtained the tracking data. I.B. discussed fish behavior and wrote the manuscript. U.G. organized and helped field work. S.B. supervised projects, wrote, and reviewed the manuscript.

## Acknowledgments

S.B. acknowledges funding support from NSF Grants CAREER iOS-1941933 and and PHY-2310691. D.C. acknowledges funding support from the National Research Foundation of Korea (NRF) grant funded by the Ministry of Science and ICT (RS-2023-00248034) and the 2024 Nerem Travel Award from the Petit Institute for Bioengineering and Bioscience at Georgia Institute of Technology. We thank the Universiti Brunei Darussalam for our Research Collaboration Approval and Agreement UBD/AVC-RI/1.21.1[a]/2024/009 and the Royal Brunei Yacht Club for supplying their resources. We thank members of the Bhamla Lab for helpful discussions, Nami Ha, Immanuel John, Joremy Tony, Katrin Grafe for their collaboration during field work, Akshay Sivakumar for preparing the equipments and Dr. Pankaj Rohilla for his valuable feedback and mentorship. Text in this paper was revised using ChatGPT-4. Animal procedures were approved by Georgia Tech IACUC (Protocol BHAMLA-A100704). The field work in South Korea (Brunei) has been conducted at 27th August 2024 (29th August 2024) and the location is 37°15’N 126°33’E (4°58’N 115°03’E).

